# Short-Term Developmental Trajectories of Dorsal-Ventral Pathways and Their Relationships with First-Grade Learning

**DOI:** 10.64898/2026.06.02.729623

**Authors:** Xueying Ren, James R. Booth, Gabriele Amorosino, Franco Pestilli, Sophia Vinci-Booher

## Abstract

The first year of formal schooling is a year of foundational reading and math learning, and individual differences emerging within this single year predict academic achievement decades later. Yet, how brain changes throughout this critical year relate to individual differences in reading and math learning remains uncharacterized. In this pre-registered study (https://osf.io/97ybe), we acquired monthly both behavioral assessments of reading- and math-learning, and diffusion-weighted MRI scans to measure white matter microstructure, across the first-grade year. Behavioral learning trajectories follow either a sigmoid for reading or an inverted-U for math. Month-to-month microstructural changes in the right middle longitudinal fasciculus predicted corresponding changes in math performance, but not in reading. Findings highlight white matter microstructure as a dynamic substrate of early math learning, and reveal a more general principle: rapid changes in white-matter microstructure during the foundational learning window may be associated with distinct academic domains.

**Funding:** R01 HD114489

## Introduction

The first year of formal schooling is a critical year when children learn basic reading and math skills that set the foundation for learning in later years (Ehri, 2005; Geary, 2011). Individual differences emerging across this single year predict academic achievement many years later (Borchers et al., 2019; Duncan et al., 2007a), yet we lack a brain-based explanation for how learning in this foundational year translates to long-term outcomes. One theory is that brain structures link foundational learning with long-term outcomes; learning in early childhood remodels brain structures that increasingly begin to facilitate but also constrain future learning (Greenough et al., 1987; Knudsen, 2004). Major white matter tracts are commonly investigated to test this theory because they connect functional brain areas, and their tissue properties correlate with performance across multiple domains (Johansen-Berg, 2010; Kanai & Rees, 2011). Indeed, research in older children during an intensive academic intervention has revealed that the tissue properties of some major white matter tracts change dynamically with performance while others remain stable and consistently predict subsequent behavioral changes (Huber et al., 2018). However, it is unknown if similar results would be obtained in younger, first grade children at the very onset of formal reading and math instruction. Here, we address this gap in knowledge by using dense neuroimaging (Vinci-Booher et al., 2025) and tracking white matter tracts tissue properties and children’s reading- and math-performance on a month-to-month basis throughout first grade. This fine-grained temporal resolution allows us to capture structural remodeling as it unfolds in this foundational year and provide insight into a potential neural pathway linking early learning to long-term outcomes.

### Foundational learning in reading and math during first grade

First grade is a critical period in academic learning with substantial changes in basic reading and math performance. For reading, most children enter first grade learning simple letter–sound correspondences and end the year learning early decoding - sounding out words, recognizing common sight words, and beginning to read simple connected text independently (Ehri, 2005). Children undergo a parallel transition in math, progressing from informal counting strategies toward symbolic number processing and basic single-digit addition and subtraction (Geary, 2004). The basic reading and math skills learned in this year are foundational in a literal sense: each subsequent year of academic learning builds upon their consolidation, and individual differences emerging during this single year are predictive of academic achievement years later (Duncan et al., 2007a).

Both reading and math learning in the first-grade are highly individualized and rarely linear (Siegler & Ellis, 1996). Reading trajectories vary markedly across children: some show rapid take-off in decoding while others lag, and divergence between typical readers and those later identified with dyslexia becomes detectable during this year (Pfost et al., 2014). Math trajectories show similar heterogeneity, with first-grade arithmetic following cumulative or compensatory patterns shaped by entering number sense, and early divergence from typical development marking children who will be identified with dyscalculia (Salaschek et al., 2014). Reading and math difficulties co-occur often - approximately 40% of children with one type of learning difficulty also experience the other (Landerl & Moll, 2010; Ren & Libertus, 2023; Willcutt et al., 2013), raising the possibility that shared white-matter pathways may underlie individual differences in both academic domains. Together, this means that group-averaged trajectories can obscure individual variability that may signal emerging disorders before behavioral differences become apparent.

### White matter and foundational learning

White-matter tracts are a leading candidate for the brain structures that link foundational learning with long-term outcomes. They provide the scaffold for inter-regional communication thought to support complex cognitive processes, and evidence indicates that their microstructure is both remodeled by early learning and, in turn, facilitates (and potentially constrains) the future learning (Wandell & Yeatman, 2013). Cross-sectional studies in school-age children (covering kindergarten through highschool) consistently demonstrate a relationship between white matter microstructure and academic learning. Both reading and math performance correlate with overlapping tracts along the left perisylvian network, including the arcuate (Arc), inferior longitudinal fasciculus (ILF), and superior longitudinal fasciculus (SLF) (Deutsch et al., 2005; Klingberg et al., 2000; Tsang et al., 2009; Wang et al., 2017), consistent with the comorbidity of dyslexia and dyscalculia (Landerl & Moll, 2010) and suggesting partially shared white-matter substrates across the two domains. However, evidence for domain-selective sub-bundles within these major tracts has also been reported (Grotheer et al., 2019). The direction of this relationship may depend on timing: learning may remodel white matter microstructure early on, while microstructure may predict future learning later.

Longitudinal and intervention studies demonstrate that reading abilities and white-matter development are dynamically linked, with reading experience remodeling white-matter structure and early white-amtter trajectories facilitating later reading outcomes. For example, in 7-to15-year-olds tracked over three years, above- and below-average readers showed divergent developmental trajectories in the left arcuate fasciculus and left inferior longitudinal fasciculus (ILF): above-average readers had lower initial FA that increased over time, while below-average readers had higher initial FA that declined over time, such that individual rates of FA development were positively correlated with reading ability (Yeatman, Dougherty, Ben-Shachar, et al., 2012). Several lines of evidence suggest that reading experience can remodel white-matter structure: in children tracked from 1st through 4th grade (mean age 7.45 at baseline), within-individual reading improvements predicted subsequent left-arcuate microstructural change rather than the reverse (Roy et al., 2025). Consistent with this, short-term reading intervention in 1st- and 2nd-graders with reading disabilities (Meisler et al., 2024) and in 7- to 12-year-olds with reading difficulties (Huber et al., 2018), produce rapid, widespread microstructural change within weeks - particularly in the left arcuate and ILF - that scales with behavioral gain. Other evidence suggests that early white-matter trajectories may facilitate later reading outcomes: Turesky et al. (2025) followed 137 children with approximately 3 diffusion MRI scans from infancy through second grade and found that the rate of left-hemisphere white matter development predicted preschool phonological processing, which in turn mediated decoding and word-reading outcomes in early elementary school.

For learning math, longitudinal evidence is sparser but implicates a partially distinct set of white-matter tracts centered on the SLF and left Arc. Evidence that math learning can remodel white matter structure comes from an 8-week intensive math intervention in 3rd graders (mean age 8.61 years) that produced changes in the SLF and Arc that scaled with individual differences in math performance increases (Jolles, Wassermann, et al., 2016). Evidence that white matter properties may facilitate math performance comes from prospective studies in which individual differences in left parietal white matter prospectively predicted math achievement in school-age children (7-9 years) (van Eimeren et al., 2008). Notably, in a longitudinal sample of 1st- through 4th-graders, increases in math performance did not track development of the left Arc (Roy et al., 2025), indicating that the left Arc mostly directly linked to reading may not be the primary substrate for math learning. Other evidence suggests that math instead may involve right-hemispheric pathways such as the right MdLF (Buianova et al., 2025) and bilateral fronto-parietal connections in the SLF (Polspoel et al., 2019).

Capturing the dynamic relations between white matter and learning requires neuroimaging designs that match the temporal resolution at which learning itself unfolds. Dense longitudinal neuroimaging (DLN), which is defined as sampling densely across a short time window, can precisely estimate individual trajectories of brain change. Currently, only a handful of studies have used a dense longitudinal approach to investigate rapid learning-related white matter changes, coming from short-term intervention contexts in struggling readers (Huber et al., 2018, 2021; Meisler et al., 2024) or older elementary-age children (Jolles, Wassermann, et al., 2016); whether the same dynamic coupling characterizes the natural unfolding of foundational first-grade learning, when most children first acquire decoding and basic arithmetic outside of intervention contexts, remains uncharacterized. Yet filling this gap is critical because individual differences emerging during this fundamental year may impact long-term learning outcomes (Duncan et al., 2007b).

### Posterior vertical pathway

Communication supporting reading and math were traditionally thought to be conveyed through the Arc, ILF, and SLF (Catani et al., 2005; Grotheer et al., 2019; Matejko & Ansari, 2015; Vandermosten et al., 2012), and the developmental trajectories and academic-learning associations of these tracts have been extensively characterized (Catani et al., 2005; Grotheer et al., 2019; Lebel et al., 2008; Matejko & Ansari, 2015; Vandermosten et al., 2012; Yeatman, Dougherty, Ben-Shachar, et al., 2012). While all three tracts are well-established for reading, math has been linked more directly to the Arc and SLF via their frontoparietal connections supporting magnitude representation and arithmetic fact retrieval (Matejko & Ansari, 2015).

More recently, a set of posterior vertical white matter tracts has come under investigation for their potential role in reading and math performance (Bullock et al., 2022a; Hayashi et al., 2024; Takemura et al., 2019; Vinci-Booher et al., 2023), owing to both advances in diffusion tractography modelling and tools that enable the identification of vertically oriented tracts (Hayashi et al., 2024; Tournier et al., 2007). The posterior vertical pathway (PVP) is well-positioned anatomically to support both reading and math learning as it directly connects temporal and parietal cortices (Bullock et al., 2022b; Takemura et al., 2019). The PVP includes two branches of the middle longitudinal fasciculus (MdLF-Ang and -SPL), the temporo-parietal connection (TPC), and the posterior arcuate fasciculus (pArc), each linking ventral temporal regions implicated in word and symbolic number processing with parietal regions implicated in attention, semantic integration, magnitude processing, and phonological processing (Bruckert et al., 2025; Bullock et al., 2019; Koirala et al., 2021; Makris et al., 2013; Weiner et al., 2017; Yeatman et al., 2011).

The existing literature relating PVP tracts with academic learning suggests that tracts within the PVP may have distinct relationships with reading and math: the posterior temporal portion of the Arc (consistent with our definition of the pArc) tracks reading changes longitudinally (Roy et al., 2025). Although direct evidence linking the TPC and academic learning is lacking, the cortical endpoints of the TPC suggest that it may be the PVP tract that is best positioned to support math as it directly connects SPL associated with numerical magnitude processing (Dehaene et al., 2003; Harvey et al., 2013) and the middle-to-posterior portion of the inferior temporal lobe associated with visual processing of symbolic numbers (Hannagan et al., 2015; Shum et al., 2013). Direct evidence linking MdLF to reading or math learning in children is lacking, leaving open the question of how MdLF microstructure relates to either domain at early developmental stage.

### Microstructural specificity: beyond FA

Above we have introduced our white matter concepts more broadly referring to microstructure. More specifically, Fractional Anisotropy (FA) is the DWI-derived measure most commonly used. FA is a composite measure influenced by multiple microstructural properties, including axonal density, myelination, and the geometric organization of fiber bundles (Jelescu & Budde, 2017). Complementary diffusion models, such as Neurite Orientation Dispersion and Density Imaging (NODDI), can further decompose the diffusion signal into indices of neurite density (NDI) and orientation dispersion (ODI) to capture how coherently fibers are aligned and to allow each microstructural property to be quantified independently (Zhang et al., 2012). Cross-sectional lifespan studies suggest that rapid FA increases during the first two decades of life are driven primarily by changes in NDI, with a negligible contribution from ODI (Chang et al., 2015; Genc et al., 2017; Mah et al., 2017). Whether the same biological processes underlie the more rapid, learning-related microstructural changes that occur over months in early childhood is, however, less clear. When children in an 8-week reading intervention showed FA gains in the arcuate fasciculus, those gains did not reflect increases in myelination or in neurite density, the biological processes thought to drive normative developmental FA increases, but instead reflected changes in the tissue surrounding axons, potentially related to glial cell activity rather than axonal change itself (Huber et al., 2021). This suggests that the biology driving the relatively fast microstructure changes associated with learning may differ from the more heavily studied slow microstructure changes across the lifespan.

### The current study

Here, using a dense longitudinal neuroimaging design with an a priori focus on the PVP (Vinci-Booher et al., 2025), we tested whether month-to-month changes in white-matter microstructure predict concurrent changes in the foundational reading and math skills that children acquire across the first-grade year–examining how brain structures may be remodeled during early learning experiences. We acquired monthly diffusion MRI and monthly behavioral assessments of reading and math from nine first-grade children across a full 12-month year, covering the months before and after the first-grade academic year (**Figure 1a**). We focused on four PVP tracts selected for their anatomical position as direct bridges between temporal and parietal cortices: the pArc, TP-SPL, MDLF-Ang and MDLF-SPL (**Figure 1b**). We used time-series modeling to distinguish the contributions of initial brain structure from ongoing brain structure remodeling and learning in both reading and math (**Figure 1c**), using FA and NODDI measures to examine microstructural properties.

**Figure 1.**
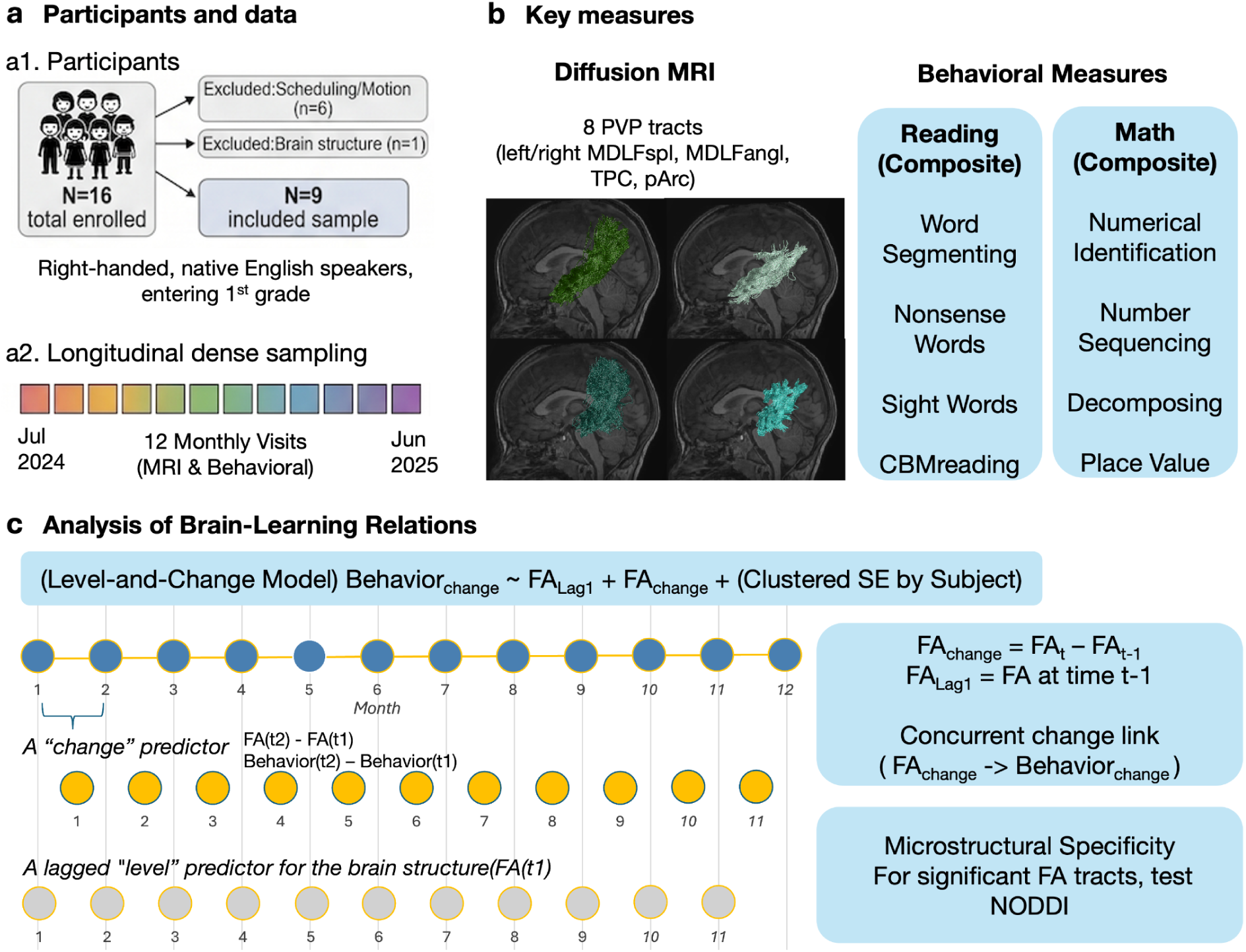
Experimental design and modeling approach. **a**. **Participant recruitment and retention.** A total of 16 participants (6-year-old) were enrolled during the summer of their first grade year. Following exclusions for scheduling confluicts, excessive head motion (*N*=6), and abnormal brain structure (*N*=1), the final analytic sample consisted of *N*=9 children. The study employed a dense longitudinal neuroimaging design consisting of 12 monthly visits over a year. Diffusion MRI and behavioral measures were acquired at each visit. **b. Key measure.** Diffusion MRI focused on 8 PVP tracts (left and right MDLFang, MDLFspl, pArc, and TPC). Behavioral performance was measured for reading and math, and composite scores were calculated for each test for later analysis. **c. Statistical modeling of brain–learning relations.** We used a Level-and-Change Model to estimate the association between white matter microstructure and monthly changes in reading and math performance. The model regressed the month-to-month change in behavioral performance on the baseline level of white matter microstructure from the previous month and the concurrent change in microstructure. Models were fitted using Ordinary Least Squares (OLS) regression with standard errors clustered by subject to account for the longitudinal nature of the data and within-subject dependence. For tract–behavior combinations showing significant FA-behavior associations, secondary analyses were conducted using Neurite Orientation Dispersion and Density Imaging (NODDI) metrics—specifically Neurite Density Index (NDI) and Orientation Dispersion Index (ODI)—to further characterize the underlying microstructural changes.

Our pre-registered predictions (https://osf.io/97ybe) were that (1) PVP tracts would show significant microstructural maturation over the first-grade year; (2) microstructural properties at each visit would predict subsequent monthly changes in reading and math on a month-to-month basis because all first-grade children in our study were required to have attended kindergarten and were expected to have some prior experience with reading and math; and (3) there would be a double dissociation among PVP tracts, such that different tracts would predict reading versus math learning. For instance, it is possible that the pArc, a key component of the language stream (Catani et al., 2005; Grotheer et al., 2019), would be significantly more predictive of reading than math (Hannagan et al., 2015). Similarly, it is possible that the TP-SPL, connecting temporal and parietal regions involved in numerical cognition and spatial processing (Bullock et al., 2019; Dehaene et al., 2003), would be significantly more predictive of math than reading.

In addition to our pre-registered analyses, we performed post hoc analyses that further interrogated the data to provide critical context for interpreting the results of our pre-registered analysis plan. Results from analyses that were not pre-registered are indicated by referencing the analysis as a “post hoc analysis” or in parentheses at the end of a sentence, i.e., (post hoc).

## Results

### Reading and math trajectories were nonlinear in distinct ways

To characterize developmental trajectories over the first-grade year, and to test whether there was consistent development at the group level, we first fitted a linear regression model to each participant’s longitudinal reading and math composite scores. Composite scores are vertically scaled with a fixed reference across all 12 sessions, so the within-child increases reported below reflect reading and math performance on a common developmental scale rather than relative-to-peer position. Then we extracted individual linear slopes for both the reading and math scores and tested whether the mean of the individual slopes was significantly different from zero.

**Table 1.**
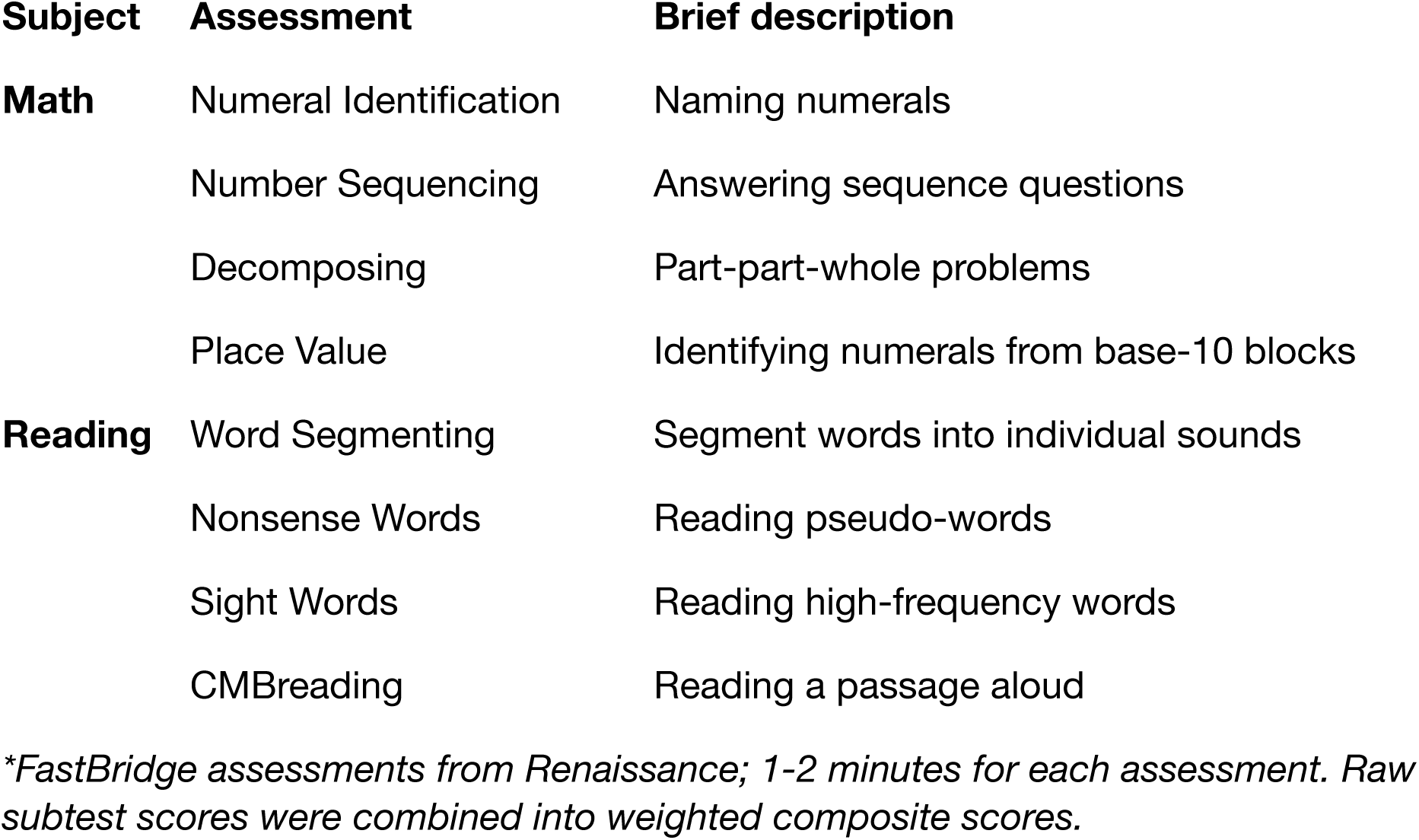
Behavioral measures for dense sampling throughout the first grade.

### Reading performance followed a sigmoid trajectory

Participants demonstrated improvements in reading, yielding a mean linear slope significantly greater than zero (mean slope = 2.15, *SD* = 0.75, *t(*8) = 8.16, *p* < .001, one-sample *t*-test, *Cohen’s d* = 2.89).

While the linear slope analysis tests whether improvement is consistent across sessions, we further characterized the nature of behavioral change using two complementary approaches. First, we conducted a paired comparison that offers a more intuitive and interpretable metric of total annual change (post hoc).

Using a paired-samples *t*-tests comparing baseline (session 1) and endline (session 12) reading scores, there was a statistically significant increase in early reading composite scores from baseline (*M* = 47.74, *SD* = 14.82) to endline (*M* = 61.37, *SD* = 11.65; *t(*8) = 4.83, *p* = .001, *Cohen’s d* = 1.61), indicating improvement over the year (**Figure 2a**). Second, early reading scores were modeled using a non-linear sigmoid (S-shaped) function *y* = *base* + *L*/(1 + *exp*(− *k*(*x* − *x*_0_))) (post hoc). The baseline parameter revealed that students started at a baseline reading level (*M_base* = 38.48, *SD* = 8.74), and the reading trajectories exhibited a qualitative shift mid-year. The inflection point (x₀), which represents the period of most rapid, steepest developmental growth, occurred on average around Session 6.68 (*SD* = 0.83) (**Figure 2b**). During this rapid-change phase, students’ reading scores increased by an average maximum amplitude (*L*) of 20.41 points (*SD* = 7.11) before establishing a new, higher plateau. Notably, the steepness of this developmental jump (*k*) showed high individual variance (*M* = 9.60, *SD* = 8.14), indicating that while some students experienced a gradual S-curve, others experienced an abrupt leap in reading proficiency.

**Figure 2.**
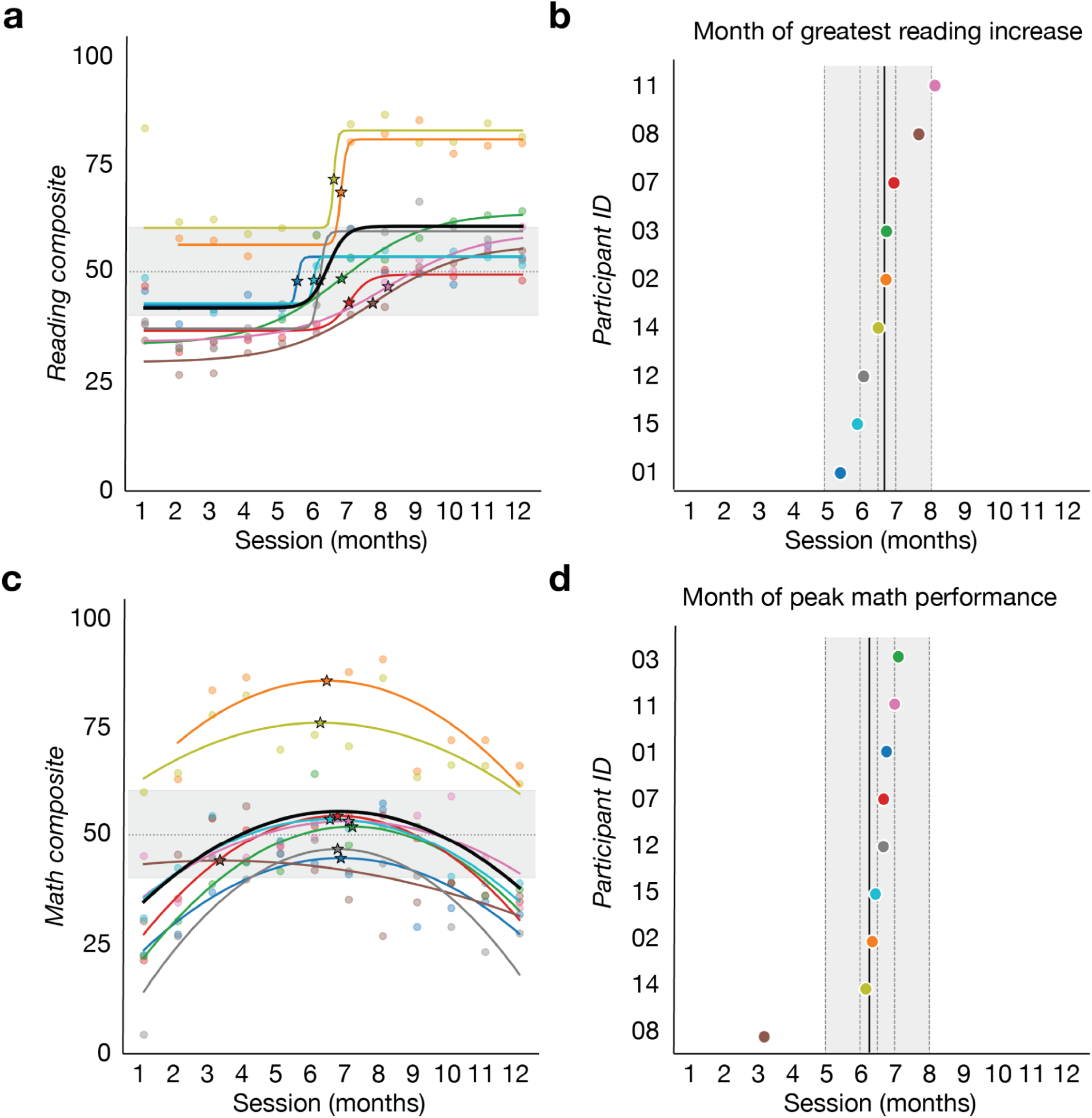
Distinct trajectories for reading and math performance across the first-grade year. Math and reading scores are reported as vertically-scaled composites (M = 50, SD = 15, anchored to the G1_Fall mean/SD of the sample and held fixed across all sessions); within-child increases reflect reading and math performance on a common developmental scale rather than movement against a shifting age norm. **a.** Individual reading composite scores across 12 monthly sessions. Thin colored lines represent individual participants (*N* = 9); the thick black line represents the group mean. **b.** Individual reading trajectories fitted with sigmoid functions (y = base + L / (1 + exp(-k(x - x₀))). The model converged for all 9 participants; for SS14, the first data point was excluded prior to fitting due to an elevated baseline. Dots represent raw data; curves represent the model fits for each participant; stars represent their inflection point (post hoc analysis). **c.** The inflection point for each subject, representing the month of greatest reading increase, occurred on average around session 6.68 (*SD* = 0.86) (post hoc analysis), indicated with a thick black line. **d.** Individual math composite scores across 12 monthly sessions. Thin colored lines represent individual participants (*N* = 9); the thick black line represents the group mean. **e.** Individual math learning trajectories fitted with quadratic functions (y = ax² + bx + c). The inverted-U pattern reflects an initial period of rapid increase followed by peak performance occurring on average around session 6.27 (*SD* = 1.18). Dots represent raw data; curves represent individual model fits; stars represent inflection points (post hoc analysis). **f.** The quadratic peak point for each subject, representing the month of peak math performance, occurred on average around session 6.27 (*SD* = 1.18) (post hoc analysis), indicated with a thick black line

Second, visual inspection of the raw trajectories suggested that linear models may not adequately capture the shape of learning across the year — reading trajectories appeared to rise rapidly before leveling off, while math trajectories showed an initial period of improvement followed by decline. Motivated by these observations, we fitted nonlinear growth models to the behavioral data: a sigmoid function for reading and a quadratic function for math (post hoc). Across the nine participants, the sigmoid model improved fit relative to the linear model (mean *R²* = 0.88 *vs.* 0.66; paired *t*(8) = 5.71, *p* < .001, one-tailed). Also, both *AIC* (Akaike Information Criterion) and *BIC* (Bayesian Information Criterion) measures favored the sigmoid on average (mean *AIC* = 70.01 *vs.* 60.61; paired *t*(8) = 4.91, *p* < .001; mean *BIC* = 70.82 *vs.* 62.22, paired *t*(8) = 4.44, *p* = .001, one-tailed). These results indicate that reading composite trajectories were better captured by a sigmoid than by a linear model.

### Math performance followed an inverted-U trajectory

Math performance did not exhibit a group-level increase. The mean linear slope for math was not significantly different from zero (mean slope = 0.05, *SD* = 0.63, *t(*8) = 0.23, *p* = .41, one-sample *t*-test, *Cohen’s d* = 0.08).

We conducted post hoc analyses, similar to the analyses for children’s reading scores. First, with paired-samples *t*-tests comparing baseline (session 1) and endline (session 12) math scores. There was a statistically significant increase in children’s early math composite scores from baseline (*M* = 33.16, *SD* = 19.11) to endline (*M* = 42.72, *SD* = 15.72), *t(*8) = 5.13, *p* = .001, *Cohen’s d* = 1.71, although this effect was not observable based on the slope of the linear fit across all 12 time points. Second, the trajectory of early math learning was modeled using a quadratic function (*y* = *ax*^2^ + *bx* + *c*) (post hoc). Math learning trajectories showed individual variability across participants. The quadratic peak (representing the inflection from rapid improvement to decline) occurred on average around session 6.27 (*SD* = 1.18), with most children peaking between sessions 6 and 7. One participant (SS08) showed a notably earlier peak around session 3, suggesting an earlier peak in math learning relative to the rest of the sample. Overall, group-level analysis of the model parameters revealed a strong initial positive linear growth rate (b; *M* = 8.39, *SD* = 3.66) combined with a negative quadratic acceleration coefficient (a; *M* = -0.64, *SD* = 0.25) (**Figure 2c & 2d**). The negative quadratic term indicates a decelerating, concave-downward trajectory; it suggests that students exhibited rapid increases in math performance during the early sessions, a peak around mid-year, and a decline in the later sessions of the first-grade year.

To verify that early math was better described by a nonlinear fit than by linear growth, we compared the linear fit against the quadratic fit as post hoc analyses. Across the nine participants, the quadratic model substantially improved fit (mean *R²* = 0.58 *vs.* 0.05; paired *t*(8) = 6.74, *p* < .001, one-tailed). *AIC* and *BIC* both favored the quadratic model on average (mean *AIC* = 85.22 *vs.* 77.07, paired *t*(8) = 4.16, *p* = .002; mean *BIC* = 86.05 *vs.* 78.31, paired *t*(8) = 3.96, *p* = .002, one-tailed). These results indicate that math trajectories were better captured by a quadratic than by a linear model.

### PVP tracts demonstrated microstructural changes over the first-grade year

Similar to our behavioral reading and math data, we first obtained a simple, interpretable summary of each child’s average rate of change to test whether the change significantly differed from zero. Specifically, we fitted individual linear models for each participant to derive a slope representing the rate of linear change in white matter microstructure (FA) for each participant in 9 tracts, including 8 tracts of interest and one comparison tract, the anterior frontal corpus callosum (ACC). To assess if the slope was significantly different from zero at the group level, we performed a one-sample *t*-test on the slope for each tract, adopting a non-parametric Wilcoxon signed-rank test for tracts that violated the assumption of normality based on a Shapiro Wilk test. Multiple comparisons were corrected using FDR (Benjamini-Hochberg) for eight tracts of interests.

The left-hemisphere tracts all exhibited mean linear slopes greater than zero, indicating FA increases over the first grade year, in the left MDLFspl (*t(*8) = 2.83, *p* = .022, *p*_FDR = .059), left MDLFang (*t(*8) = 2.90, *p* = .020, *p*_FDR = .059), left TPC (*t(*8) = 3.25, *p* = .012, *p*_FDR = .059), and left pArc (*t(*8) = 2.40, *p* = .043, *p*_FDR = .086), though they are trending at the significance level after FDR correction. In contrast, none of the right-hemisphere tracts exhibited mean linear slopes that were greater than zero, right MDLFspl (*t(*8) = 1.08, *p* = .310), right MDLFang (*t(*8) = 1.31, *p* = .227), right TPC (*t(*8) = 1.93, *p* = .089), and right pArc (*t(*8) = 2.16, *p* = .063). The ACC comparison tract also showed no change (*t(*8) = -0.31, *p* = .761). Lastly, FA in the ACC comparison tract also showed no change across the learning period (*p* = .864).

We conducted a post hoc analysis with paired-samples *t*-tests comparing FA at baseline (session 1) and endline (session 12) for each tract of interest (**Figure 3**). Multiple comparisons were corrected using FDR for eight tracts of interests. Consistent with the pre-registered analyses indicating mean linear slopes greater than zero at the group level, the post hoc analyses revealed a significant increase in the left MDLFspl FA from baseline (*M* = 0.38, *SD* = 0.03) to endline (*M* = 0.40, *SD* = 0.03), *t(*8) = 15.19, *p* < .001, *p*_FDR < .001, *Cohen’s d* = 5.06, and the left MDLFang FA from baseline (*M* = 0.34, *SD* = 0.02) to endline (*M* = 0.36, *SD* = 0.02), *t*(8) = 4.62, *p* = .002, *p*_FDR = .007, *Cohen’s d* = 1.54, and the left TPC FA from baseline (*M* = 0.40, *SD* = 0.03) to endline (*M* = 0.42, *SD* = 0.03), *t(*8) = 2.94, *p* = .019, *p*_FDR = .038, *Cohen’s d* = 0.98, and the left pArc FA from baseline (*M* = 0.41, *SD* = 0.02) to endline (*M* = 0.43, *SD* = 0.03), *t*(8) = 2.99, *p* = .018, *p*_FDR = .038, *Cohen’s d* = 1. There was no significant baseline to endline differences in FA for any right hemisphere tract (all *ps* > 0.10) except for the right TPC FA that increased from baseline (*M* = 0.39, *SD* = 0.02) to endline (*M* = 0.40, *SD* = 0.02), *t*(8) = 2.57, *p* = .033, *p*_FDR = .053, *Cohen’s d* = 0.86, but trending at significance level after multiple comparison correction. All post-hoc analyses were consistent with the pre-registered analyses, revealing microstructural changes across the first grade year for all PVP tracts except right MDLFspl, right MDLFang, and right pArc.

**Figure 3.**
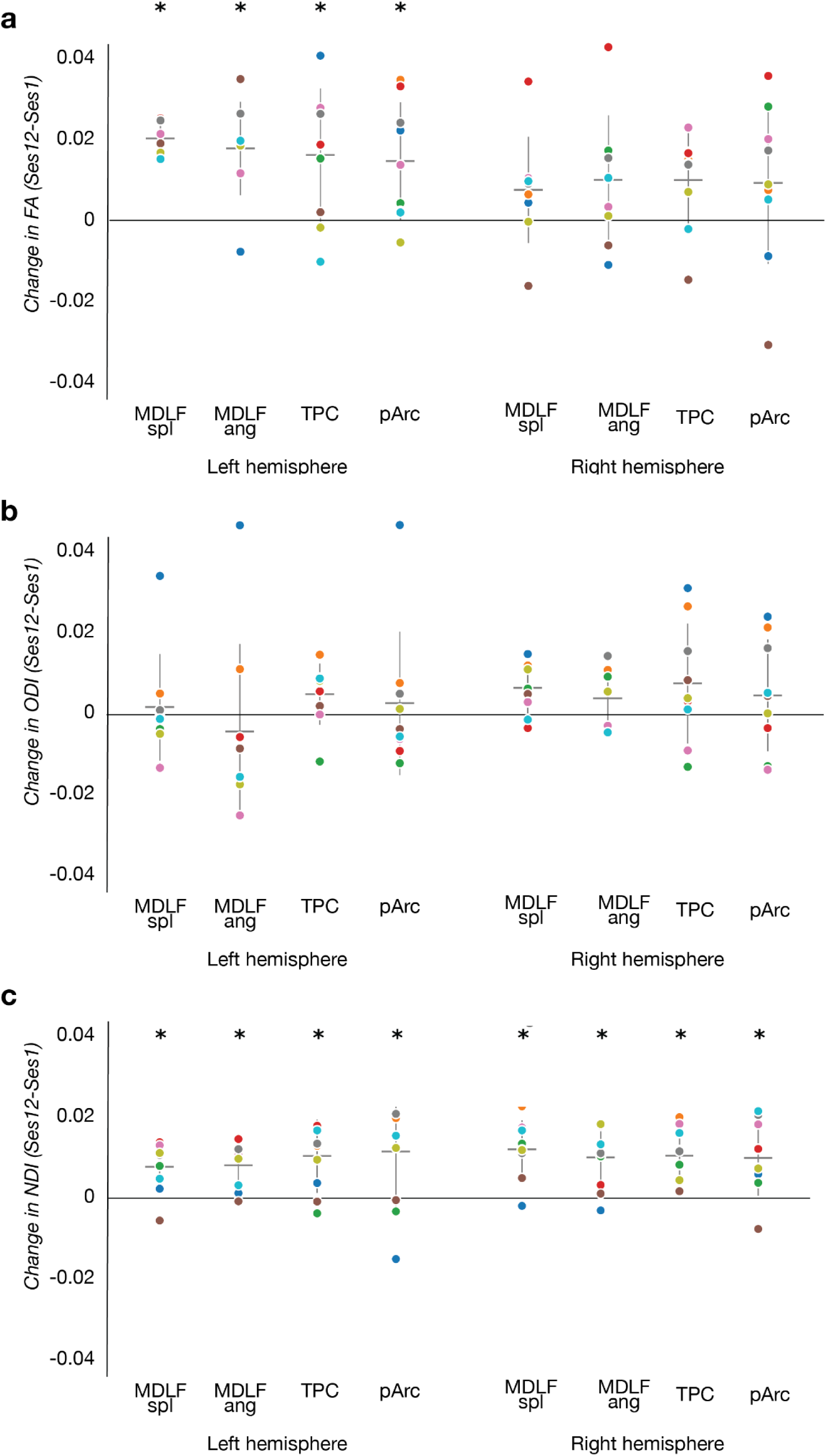
PVP tracts demonstrated microstructure changes across the first-grade year. **a.** Fractional anisotropy (FA). **b.** Orientation dispersion index (ODI). **c.** Neurite density index (NDI). Dot plots following each panel represent FA change from baseline (session 1) to endline (session 12) for each individual (*N* = 9), and the asterisks indicate significance at the group level after FDR correction for 8 tracts. Tracts shown include the bilateral posterior vertical pathway (PVP) tracts — middle longitudinal fasciculus (MDLFspl, MDLFang), temporo-parietal connection (TPC), and posterior arcuate (pArc). Colors are consistent across all panels and match those used in subsequent figures to reference individual participants.

Finding no baseline-to-endline changes in FA in the right PVP tracts (right MDLFspl, right MDLFang, and right pArc) was unexpected, and it remained possible that the underlying microstructural metrics, NDI and ODI, were changing in ways that were not reflected in FA (Zhang et al., 2012). To test this, we conducted a post hoc analysis with paired-samples *t*-tests comparing NDI and ODI at baseline (session 1) and endline (session 12) for all PVP tracts. Multiple comparisons were corrected using FDR for eight tracts (**Figure 3**).

For the neurite density (NDI), the NODDI follow-up analyses revealed a significant increase in the rightMDLFspl NDI from baseline (*M* = 0.45, *SD* = 0.02) to endline (*M* = 0.46, *SD* = 0.02), *t*(8) = 5.05, *p* = .001, *p*_FDR = .005, *Cohen’s d* = 1.68, the rightMDLFang NDI from baseline (*M* = 0.45, *SD* = 0.02) to endline (*M* = 0.46, *SD* = 0.01), *t*(8) = 3.81, *p* = .005, *p*_FDR = .009, *Cohen’s d* = 1.27, the rightTPC NDI from baseline (*M* = 0.48, *SD* = 0.02) to endline (*M* = 0.49, *SD* = 0.02), *t*(8) = 4.58, *p* = .002, *p*_FDR = .005, *Cohen’s d* = 1.53, the rightpArc NDI from baseline (*M* = 0.48, *SD* = 0.02) to endline (*M* = 0.49, *SD* = 0.02), *t*(8) = 3.20, *p* = .013, *p*_FDR = .014, *Cohen’s d* = 1.07, the leftMDLFspl NDI from baseline (*M* = 0.46, *SD* = 0.02) to endline (*M* = 0.47, *SD* = 0.02), *t*(8) = 3.73, *p* = .006, *p*_FDR = .009, *Cohen’s d* = 1.24, the leftMDLFang NDI from baseline (*M* = 0.45, *SD* = 0.02) to endline (*M* = 0.46, *SD* = 0.02), *t*(8) = 4.50, *p* = .002, *p*_FDR = .005, *Cohen’s d* = 1.50, the leftTPC NDI from baseline (*M* = 0.49, *SD* = 0.02) to endline (*M* = 0.50, *SD* = 0.02), *t*(8) = 3.44, *p* = .009, *p*_FDR = .012, *Cohen’s d* = 1.15, and the leftpArc NDI from baseline (*M* = 0.50, *SD* = 0.03) to endline (*M* = 0.51, *SD* = 0.02), *t*(8) = 2.36, *p* = .046, *p*_FDR = .046, *Cohen’s d* = 0.79. For the orientation dispersion (ODI), the same analyses revealed an increase in the rightMDLFspl ODI from baseline (*M* = 0.21, *SD* = 0.01) to endline (*M* = 0.21, *SD* = 0.01), *t*(8) = 3.16, *p* = .013, *p*_FDR = .107 *Cohen’s d* = 1.05, but did not pass the FDR correction. ODI did not show significant increase in other PVP tracts (*ps* > .08).

As post hoc analyses, to link our findings with prior literature (Broce et al., 2019; Grotheer et al., 2019), we also examined the vertical occipital fasciculus (VOF), a vertical posterior pathway adjacent to the PVP tracts (Bullock et al., 2019; Vinci-Booher et al., 2022; Yeatman et al., 2014), as well as the Arcuate (Arc) and the Superior Longitudinal Fasciculus (segmented as SLF1and2 and SLF3) that have been associated with reading and math learning in prior work (Grotheer et al., 2019; Klingberg et al., 2000; Tsang et al., 2009; Yeatman, Dougherty, Ben-Shachar, et al., 2012) using a paired-samples *t*-tests comparing FA at baseline (session 1) and endline (session 12). Multiple comparison were corrected using FDR for those either tracts. The post hoc analyses revealed a significant increase before multiple comparison correction, but the results were trending after the correction. Specifically, there is an increase in the leftArc FA from baseline (*M* = 0.39, *SD* = 0.02) to endline (*M* = 0.40, *SD* = 0.02), *t*(8) = 2.88, *p* = .020, *p*_FDR = .082, *Cohen’s d* = 0.96, the leftSLF1&2 FA from baseline (*M* = 0.35, *SD* = 0.01) to endline (*M* = 0.36, *SD* = 0.02), *t*(8) = 2.55, *p* = .034, *p*_FDR = .089, *Cohen’s d* = 0.85, the leftVOF FA from baseline (*M* = 0.32, *SD* = 0.03) to endline (*M* = 0.33, *SD* = 0.03), *t*(8) = 2.38, *p* = .044, *p*_FDR = .089, *Cohen’s d* = 0.79, and the rightVOF FA from baseline (*M* = 0.32, *SD* = 0.02) to endline (*M* = 0.34, *SD* = 0.02), *t*(8) = 3.63, *p* = .007, *p*_FDR = .053, *Cohen’s d* = 1.21. In contrast, no significant change in FA was observed in the right Arc, rightSLF1&2, leftSLF3, or the rightSLF3 (all *ps* > .10).

### Month-to-month behavioral changes relate to month-to-month changes in tract microstructure

To test whether the microstructural properties would prospectively predict subsequent monthly changes in reading and math on a month-to-month basis, we adopted a complementary approach (pre-registered analysis with statistically necessary deviations) to the preregistered analysis. Specifically, a “level-and-change” model (defined below) was used to test whether prior white matter measurements or concurrent white matter changes predicted changes in reading or math performance (pre-registered analysis with deviation of models; Methods Statistical analysis section). The “level-and-change” model regresses changes in behavioral scores on baseline FA (FA level at t-1) and month-to-month FA change (moving FA change, mΔFA hereafter) using OLS regression with standard errors clustered by subject, as linear mixed-effects models failed to converge. To control for four multiple comparisons, FDR correction was applied.

### MDLF microstructure predicts concurrent change in math performance

Results demonstrated that month-to-month changes in FA of right MDLFspl and MDLFang tracts predicted month-to-month changes in math performance. The right MDLFspl and right MDLFang both showed significant associations between mΔFA and math score changes after FDR correction. Specifically, a 0.01 unit increase in session-to-session FA was associated with a 1.04 point increase in the right MDLFspl (right MDLFspl: *β* = 103.61, *t* = 2.26, *p* = .024, *p*_FDR = .047; model fit: *F*(2, 85) = 4.47, *p* = .050, *R²* = .019), and with a 1.10 point increase in math score change in the right MDLFang (right MDLFang: *β* = 109.70, *t* = 2.29, *p* = .022, *p*_FDR = .047; model fit: *F*(2, 85) = 2.91, *p* = .112, *R²* = .030). These positive coefficients indicate that sessions in which FA increased in the right MDLF were associated with increased math changes (**Figure 4**). In both tracts, the baseline FA level (FA_Lag1) did not significantly predict math performance change (right MDLFspl: *β* = 22.67, *t* = 0.75, *p* = .456; right MDLFang: *β* = -4.23, *t* = -0.16, *p* = .875), indicating that the brain-behavior coupling was associated with concurrent month-to-month FA change rather than baseline white matter structure. There were no significant relationships between FA in other PVP tracts with math after FDR correction (*ps* > .09). We also found no evidence for a relationship between FA in any PVP tracts and reading performance (all *ps* > .06).

**Figure 4.**
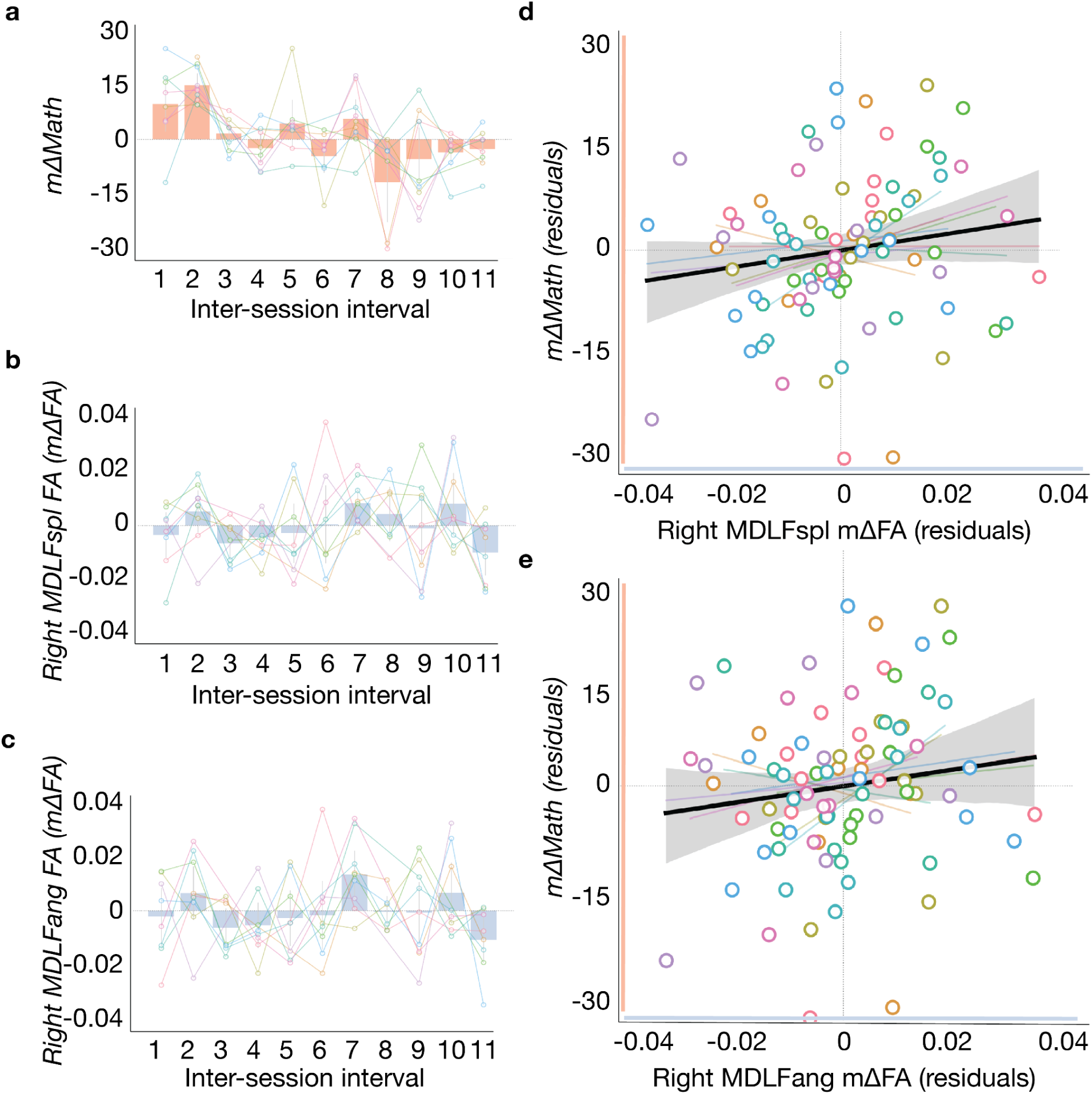
Right MDLF FA changes track concurrent month-to-month math changes. **a.** Session-to-session changes in math composite scores (mΔMath). **b**. Session-to-session changes in fractional anisotropy (mΔFA) across 11 monthly sessions in right MDLFspl. Panel **c.** Session-to-session changes in fractional anisotropy (mΔFA) across 11 monthly sessions in right MDLFang. Panel a-c, bars represent group means *(N* = 9), error bars represent 95% confidence intervals (1.96 × SEM), and colored lines connect individual participants across sessions, with white-filled dots indicating individual data points. **d.** Visualization of the brain-behavior relationship for the right MDLFspl tract and math learning. **e.** Visualization of the brain-behavior relationship for the right MDLFang tract and math learning. Axes show residuals of mΔFA and mΔMath after regressing each on baseline FA (FA_lag1); the slope equals the partial regression of mΔMath on mΔFA controlling for baseline FA (FA at t-1).. White-filled dots represent individual participant-session observations, colored by participant. Faint colored lines show per-participant best-fit lines; the black line indicates

We also tested other tracts, including the VOF, Arc, SLF1and2, and SLF3. These tracts were selected for post hoc analyses because they have been associated with reading and math learning in prior work (Grotheer et al., 2019; Matejko & Ansari, 2015; Vandermosten et al., 2012). We conducted post hoc analyses to better link our findings with prior literature by applying the level-and-change approach to FA in the VOF, Arc, SLF1and2, and SLF3 that have been associated with reading and math learning (Ben-Shachar et al., 2007; Grotheer et al., 2019; Klingberg et al., 2000; Tsang et al., 2009; Yeatman, Dougherty). In the left Arc, FA change significantly predicted math change (*β* = -123.15, *t* = -3.02, *p* = .003, *p*_FDR = .023; Model fit: *F*(2, 85) = 4.85, *p* = .042, *R²* = .034), but not reading (*p* = .26). The negative coefficient indicates that sessions with greater FA increases in the left Arc were associated with smaller concurrent math gains, an effect opposite in direction to that observed in the right MDLF tracts. Baseline FA level did not significantly predict math change in the left Arc (*p* = .569). We used the same level-and-change framework for other additional tracts, however, we did not find significant relationships between FA in either tracts with math or reading (all *ps* > .15). The comparison tract, anterior frontal corpus callosum, also did not show significant associations with either math or reading (*ps* > .06).

### Changes in orientation dispersion, not neurite density, predict math learning

To further investigate the microstructural changes underlying the FA change, we decomposed the FA into indices of neurite density (NDI) and orientation dispersion (ODI) using the NODDI method (Zhang et al., 2012). Using the same level-and-change OLS framework with standard errors clustered by subject, we tested whether month-to-month changes in neurite density index (NDI) or orientation dispersion index (ODI) predicted math score changes for the MDLF tracts identified in the prior FA analyses. Because these analyses targeted tracts that had already survived FDR correction in the primary FA analysis, raw *p*-values are reported.

Changes in ODI (mΔODI), but not mΔNDI, significantly predicted math score changes in both right MDLF tracts. In the right MDLFspl, month-to-month decreases in orientation dispersion were associated with greater math performance changes (*β* = -237.18, *t* = -2.12, *p* = .034; **Figure 5**), and a similar pattern was observed in the right MDLFang (*β* = -278.84, *t* = -2.43, *p* = .015). ODI level at the previous month was also not a significant predictor (right MDLFang: *p* = .625; right MDLFspl: *p* = .117). Neither mΔNDI nor NDI level significantly predicted mΔMath in either tract (right MDLFang: *p* = .283; right MDLFspl: *p* = .242). These results suggest that the FA–math relationship is driven by increasing fiber coherence — that is, white matter fibers becoming more uniformly oriented — rather than by changes in neurite density. This pattern is consistent with a maturing white matter architecture in which the organizational refinement of existing fiber populations, rather than the proliferation of new neurites, supports the learning of math during the first year of formal schooling.

**Figure 5.**
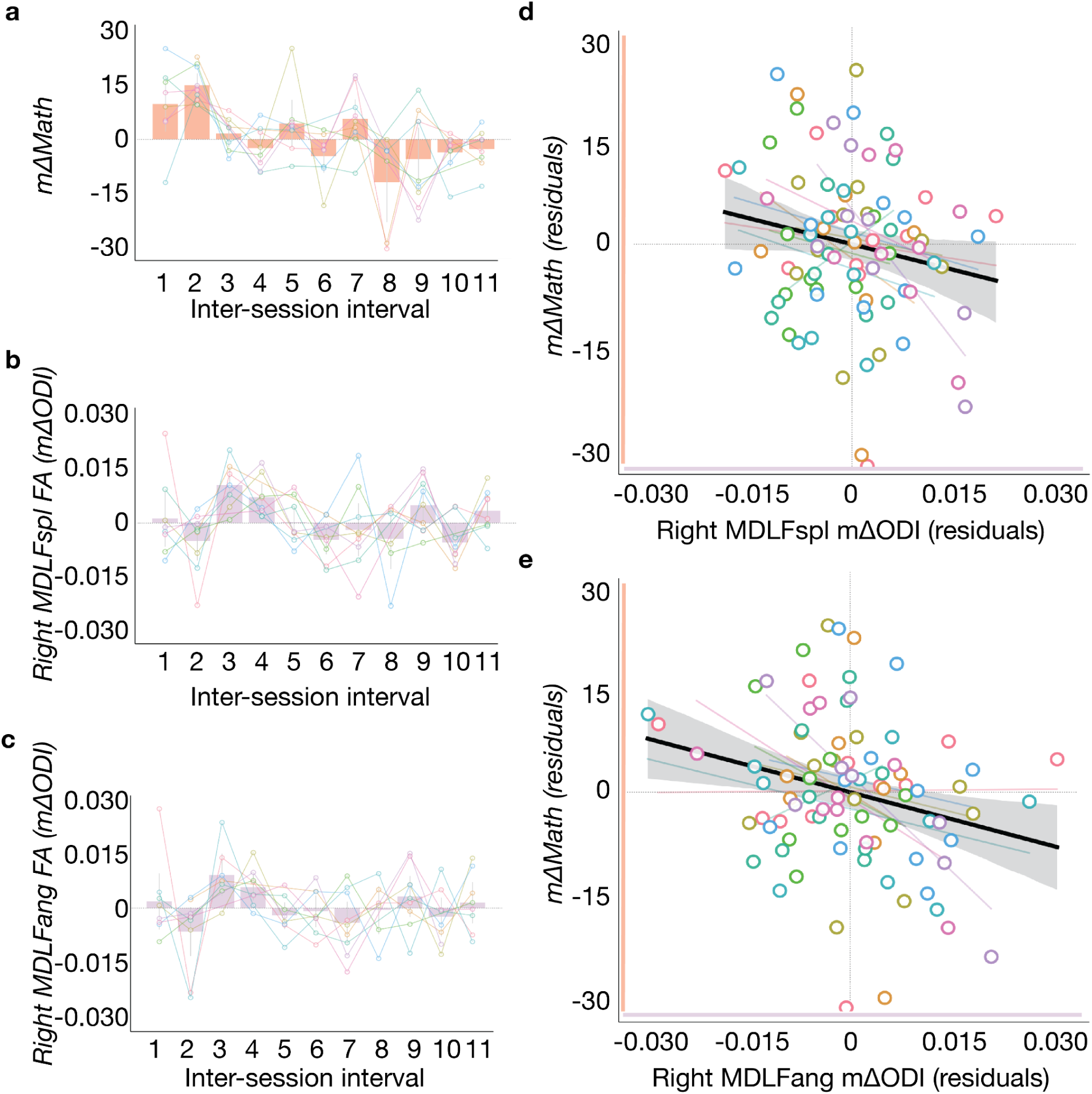
**Changes in orientation dispersion tack changes in math performance (pre-registered exploratory analysis with necessary deviation of models)**. **a.** Session-to-session changes in math composite scores (mΔMath). **b.** Session-to-session changes in orientation dispersion index (mΔODI) across 11 monthly inter-session intervals in right MDLFspl. **c.** Session-to-session changes in orientation dispersion index (mΔODI) across 11 monthly inter-session intervals in right MDLFang. In panels a-c, bars represent group means (N = 9), error bars represent 95% confidence intervals (1.96 × SEM), and colored lines connect individual participants across sessions, with white-filled dots indicating individual data points. **d.** Visualization of the brain-behavior relationship between right MDLFspl orientation dispersion and math learning. **e.** Visualization of the brain-behavior relationship between right MDLFang orientation dispersion and math learning. Axes show residuals of mΔODI and mΔMath after regressing each on baseline ODI (ODI at t-1); the slope equals the partial regression of mΔMath on mΔODI controlling for baseline ODI. White-filled dots represent individual

To probe the microstructural basis of the left Arc–math coupling, we conducted NODDI follow-up analyses using the same level-and-change framework (post hoc). Neither neurite density (NDI) nor orientation dispersion (ODI) significantly predicted mΔMath in the left Arc (*ps* > .43). Unlike the right MDLF tracts, where ODI changes paralleled the FA effect, the FA–math coupling in the left Arc was not accompanied by a corresponding signal in either NODDI metric.

### The right MDLF FA–math coupling is distributed across the year rather than localized to a critical window

To determine whether specific periods of white matter change were more predictive of overall math score changes than others, we conducted a critical windows analysis on the tracts that showed significant FA–math associations (pre-registered analysis with statistically necessary deviation of models). For each tract, FA change was decomposed into three non-overlapping windows: initial (sessions 1–3), ongoing (sessions 3–10), and recent (sessions 10–12) to reflect sparser sampling with a baseline sample (session 1), a fall semester sample (session 3), and spring semester sample (session 10), and an endline sample (session 12) as is commonly done in other works (Huber et al., 2018). These windows were entered as predictors of total math score change (session 12 minus session 1) using OLS regression with standard errors clustered by subject. A comprehensive “horse race” model including all three windows simultaneously was also tested. Because these analyses targeted tracts that had already survived FDR correction, raw *p*-values are reported.

For the right MDLFang (N = 9 with complete data across all required time points): No individual window significantly predicted total math score change. In the single-window models, the initial window showed the largest (but non-significant) coefficient (*β* = 61.73, *t* = 0.77, *p* = .441), while the ongoing (*β* = 36.01, *t* = 0.15, *p* = .884) and recent (*β* = -30.54, *t* = -0.21, *p* = .834) windows were also non-significant. The comprehensive model confirmed this pattern, with no window emerging as a significant predictor (initial: *p* = .710; ongoing: *p* = .906; recent: *p* = .937). For right MDLFspl (N = 9), similarly, no critical window was identified. None of the single-window models reached significance (initial: *β* = -54.42, *p* = .795; ongoing: *β* = -1.32, *p* = .996; recent: *β* = 48.76, *p* = .772), and the comprehensive model showed no significant predictors (all *ps* > .88). These results suggest that the relationship between right MDLF white matter changes and math learning is not driven by a specific critical period of plasticity. Rather, the FA–math association appears to reflect a continuous, distributed coupling between white matter microstructure and learning across the entire learning period.

### Testing domain selectivity: Is the pArc associated with reading and TPC associated with math?

To answer the question of whether a given tract is selectively coupled to one cognitive domain, we used the level-and-change modeling approach to test for a selective relationship between the pArc and reading and between the TPC and math. However, neither the pArc–reading nor the TPC–math associations reached significance after FDR correction for four comparisons. In the pArc–reading analysis, left pArc FA level showed a trend toward predicting reading changes (*β* = -17.61, *t* = -1.89, *p* = .059, *p*_FDR = .118; model fit: *F*(2, 86) = 1.78, *p* = .229, *R²* = .008), and right pArc FA level was not a significant predictor (*p* = .185). Neither left nor right pArc mΔFA predicted reading changes (left: *p* = .757; right: *p* = .502). In the TPC–math analysis, right TPC mΔFA showed a trend (*β* = 62.06, *t* = 2.02, *p* = .044, *p*_FDR = .088; model fit: *F*(2, 85) = 3.13, *p* = .099, *R²* = .008) but did not survive correction, and the level predictor was not significant either (*p* = .582). Left TPC showed no significant associations with math change for either FA level (*p* = .696) or FA change (*p* = .094).

We conducted post-hoc analyses to test whether the association between FA in the right MDLF and math reflected domain-selective coupling (i.e., a single dissociation), given that we found an association between right MDLF and math rather than an association between TPC and math performance as we had expected. We fit an OLS model with an interaction term between session-to-session FA change and cognitive domain (math *vs.* reading), as we did in our pre-registered analyses. Session-to-session changes in reading and math performances were z-scored within each domain so that the interaction coefficient reflected a scale-comparable difference in coupling between session-to-session math and reading performance changes in session-to-session FA changes. A significant interaction would indicate that the tract is preferentially coupled to one domain over the other. The right MDLFang showed a trend-level interaction (*β* = 12.99, *t* = 1.73, *p* = .084, *R²* = .015), suggesting that the relationship may differ between math and reading, but this effect did not reach statistical significance. The right MDLFspl showed no evidence of domain-selective coupling (*β* = 6.17, *t* = 0.56, *p* = .573, *R²* = .010). Thus, although both right MDLF sub-tracts showed significant month-to-month coupling with math learning in the primary analysis, we did not find statistical evidence that these associations were domain-selective. This should be interpreted with caution given the limited sample size (*N* = 9) and the fact that the interaction test requires detecting a difference between two already-small effects, a particularly demanding statistical comparison.

## Discussion

The first year of formal schooling is one of the most consequential periods of human cognitive development: it is when children acquire the foundational reading and math skills that scaffold all subsequent academic learning. Furthermore, individual differences emerging within this single year predict achievement decades later (Ritchie & Bates, 2013). To date, dense longitudinal diffusion MRI studies of learning have been confined to short-term educational interventions in struggling readers and older elementary-age children (Huber et al., 2018; Jolles, Supekar, et al., 2016; Meisler et al., 2024)., leaving the foundational reading and math learning that occurs during the first-grade year uncharacterized. Here, by acquiring twelve monthly assessments of behavior and white-matter microstructure across the first-grade year, we show that brain–behavior coupling during this period operates on short timescales. Behavioral trajectories were nonlinear with distinct forms for reading (sigmoid) and math (inverted-U) that were consistent across participants but that showed between-participant variances in descriptive parameters, such as the peak of the inverted-U. White-matter microstructure changes were highly individualized, with no clear group-level trends aside from an increase in FA for four left-hemisphere PVP tracts and an increase in ODI for the right MDLF tracts that comprise the anterior portion of the right PVP. Critically, our results suggest a dynamic coupling between behavior and microstructure changes: month-to-month FA (and ODI) change in the right middle longitudinal fasciculus (MDLF) predicted concurrent month-to-month changes in math performance, suggesting that foundational math learning may be associated with changes in the cohesion of right hemisphere connections between anterior temporal and parietal cortices. Together, these results identify the anterior PVP as a dynamic substrate of early math learning and demonstrate that early learning may be accompanied by rapid changes in white matter microstructure.

### Reading and math trajectories were nonlinear in distinct ways

Children showed qualitatively distinct changes in both math and reading across the first year of formal schooling. Reading score rose across the year, with a group-level linear slope significantly greater than zero and an average 13.6-point increase from baseline to endline; math score showed no significant overall linear slope yet rose with an average 9.56-point increase from baseline to endline. Critically, based on our post hoc analyses, modeling each child’s trajectory revealed two qualitatively different shapes of development: reading was best captured by a sigmoid (S-shaped) function with a clear inflection point around Session 7 at the group level (corresponding to the first month of the spring semester), whereas math followed an inverted-U trajectory with peak performance around Session 6 at the group level (corresponding to the last month of the fall semester). One possibility is that reading and math learning differ in the cohesiveness of their content during first grade. Reading at this age is a relatively unitary endeavor: children are learning to read, with sequential skills (letter-sound correspondences, fluent word recognition) that build cumulatively toward a single integrated ability (Castles et al., 2018; Ehri, 2005). First-grade math, by contrast, comprises a series of partially distinct competencies, such as counting, addition and subtraction, and place values that develop on partially independent timelines (Geary, 2004; LeFevre et al., 2010). The content of reading learning at this age is therefore more cohesive and stable than the content of math learning, which may produce smoother, more linearly accumulating gains in reading and more uneven, multi-component trajectories in math.

These shape-level findings could not have been revealed without dense within-year sampling, and they directly motivate the value of the design we adopt here. With a conventional pre-to-post or fewer-timepoint longitudinal design, our reading data would have appeared as a roughly linear gain, masking the abrupt mid-year transition that defines first-grade reading acquisition for many children. More strikingly, our math data would have produced contradictory conclusions depending on which timepoints were sampled: a pre-to-post comparison would have shown an increase in math ability (as our baseline-to-endline *t*-test confirms), but a linear slope across all twelve sessions would have suggested no development at all. By measuring math skills frequently throughout the year, we resolved these contradictions and uncovered a more accurate description of the shape of children’s math learning: an inverted U in which children make rapid early gains as they master basic math skills (e.g., counting), then started to decline gradually after a short peak around mid-year (Adolph et al., 2008; Aunola et al., 2004; Salaschek et al., 2014). This is, to our knowledge, one of the first demonstrations of nonlinear behavioral dynamics in early elementary school.

The sigmoid form of reading growth itself converges with longitudinal evidence that early reading development is highly nonlinear and individually heterogeneous, with rapid take-off periods rather than steady linear gains (Aunola et al., 2004; Pfost et al., 2014). In our sample, the steepness of this transition (k) varied across children, with some moving through a gradual S-curve and others showing an abrupt leap in reading proficiency. The temporal location of the reading inflection point and the math peak within roughly one session of each other, near the middle of first grade, further suggests that mid-year may represent a special window of learning development across domains, a within-year critical window that, again, only becomes visible at high temporal resolution and that motivates the focused white-matter analyses presented below.

### PVP tracts demonstrated microstructural changes over the first-grade year

Across the four bilateral tracts comprising the posterior vertical pathway (PVP), MDLFspl, MDLFang, TPC, and pArc, every left-hemisphere tract showed a significant FA increase across the academic year in post hoc baseline-to-endline comparisons. By decomposing FA into NDI and ODI, we found a significant baseline-to-endline increase in NDI across the four bilateral PVP tracts, and we did not observe such patterns in ODI. These results suggest that the first year of formal schooling may be a period of measurable left-lateralized FA change in the PVP, and are consistent with prior work reporting that changes in FA measured before and after first grade are associated with underlying changes in NDI (Genc et al., 2017).

Our FA and NDI results are consistent with the broader class of non-myelinating, experience-driven white matter changes identified in prior works. Longitudinal studies measuring yearly changes have established that FA in major association tracts continues to increase into early adolescence, with the steepest gains in early childhood and a slower pace through the school-age years (Lebel et al., 2008; Lebel & Beaulieu, 2011; Reynolds et al., 2019). Recent large-scale lifespan modeling of normative brain charts of white-matter microstructure across more than 26,000 scans from 42 harmonized studies likewise places the school-age window as one of continued, tract-specific FA accrual rather than a plateau (Kim et al., 2026). However, fine-grained data on rapid white matter changes during short but critical intervals of the lifespan remain sparse. What evidence does exist suggests that FA can change rapidly with experience in this age range — an 8-week reading intervention produced widespread changes in MD and FA in school-aged children within just 2–3 weeks (Huber et al., 2018), and a follow-up study associated those rapid changes to extra-axonal diffusivity rather than axonal water fraction or T1 (Huber et al., 2021). Notably, no prior conventional or dense longitudinal study included the PVP tracts examined here in the current study. The most directly relevant prior work demonstrated that PVP microstructure is more adult-like than later-maturing fronto-parietal tracts in 5- to 8-year-olds, but is not yet at the adult endpoint (Vinci-Booher et al., 2022). Our results are consistent with that placement: FA and NDI in the PVP are changing meaningfully across the first grade year.

Finally, it is important to note that finding no significant group-level FA increases in the right hemisphere tracts across the full year does not necessarily mean these tracts are microstructurally static during first grade. FA is a composite measure that depends positively on neurite density and inversely on orientation dispersion (Zhang et al., 2012). In our data, we found significant baseline-to-endline increase in NDI across four bilateral PVP tracts, while ODI showed no significant change. This is consistent with prior work showing that developmental FA changes in early childhood are primarily driven by NDI rather than ODI (Chang et al., 2015; Genc et al., 2017). Notably, the right MDLF showed a significant NDI increase despite no detectable FA change, indicating that FA alone may not fully capture microstructural development; decomposing the FA into NODDI metrics may provide more detailed readout of the specific biophysical changes occurring.

### Right-MdLF microstructural change tracks first-grade math changes

When we modeled month-to-month score change as a function of both baseline FA (level) and concurrent FA change (mΔFA) within each tract (pre-registered analysis with statistically necessary deviations), a pattern emerged: in the right MdLF, both subdivisions showed coupling between mΔFA and mΔmath, such that intervals over which a child’s right MdLFang or right MdLFspl FA increased were also intervals over which that child’s math performance increased. No PVP tract was associated with reading, and the ACC comparison tract was associated with neither domain, suggesting that the right-MDLF–math coupling might be capturing a selective relationship between right MDLF and math rather than a general developmental change.

The right-MDLF result is anatomically and developmentally plausible. The MDLF connects anterior temporal cortex with inferior and superior parietal regions, most prominently the angular gyrus and superior parietal lobule (Kalyvas et al., 2020; Makris et al., 2013), regions implicated in number sense, spatial-numerical mappings such as the mental number line, and magnitude estimation (Dehaene et al., 2003). Recent developmental work has shown that right MDLF microstructure is preferentially associated with accuracy on easier one- and two-digit arithmetic - problems related to magnitude-based processing - with influence on math performance that is most pronounced during childhood (10-13 years old) and diminishes with age (Buianova et al., 2025), a profile that maps closely onto the first-grade math performance captured by our study. The convergence between Buianova et al.’s cross-sectional right-MdLF–easier-math association and our longitudinal right-MdLF–ΔFA–math coupling lends mutual support to a right-lateralized role for the MdLF in early math learning. Our findings of brain-learning change aligns with the experience-driven framing emerging from intervention studies and longitudinal designs in older children, which find that, in childhood, learning shapes white matter rather than the reverse (Huber et al., 2018, 2021; Roy et al., 2025).

A natural question raised by these results is why we did not find a comparable brain–behavior coupling for reading, given the well-established association between the pArc and reading in older children (Yeatman et al., 2011; Roy et al., 2025). Notably, we also tested reading-related tracts (left arcuate, SLF), and did not find brain-behavior coupling with reading, indicating that the absence of a reading effect is not specific to the PVP tracts but reflect a broader characteristic of first-grade reading development at the temporal scale we sampled. It is possible that the PVP’s role in reading may emerge later in elementary school, once children move beyond decoding to engage in more semantic and orthographic processing supported by temporo-parietal connectivity (Wandell & Yeatman, 2013). Second, our dense monthly sampling within first grade allowed us to directly test whether the year-to-year pArc-reading coupling reported by Roy et al., 2025 also manifests at finer temporal scales; the absence of such coupling here suggests that the experience-driven remodeling of the pArc may accumulate gradually over multiple years rather than emerging within a single academic year.

### Microstructural basis of the right-MdLF–math coupling: orientation refinement, not neurite proliferation

To probe the biophysical changes underlying the FA–math coupling in the right MdLF, we decomposed the FA measure into its NODDI components (pre-registered analysis with statistically necessary deviations). Across both right-MdLF subtracts, concurrent changes in orientation dispersion index (ODI), but not in neurite density index (NDI), predicted month-to-month math changes, and the direction of the ODI-math relationship was negative: sessions in which a child’s right MdLF fibers became more coherently aligned (lower ODI) were the same sessions in which that child showed the largest increases in math performance. The FA–behavior coupling identified in Section 4.3 is therefore better described as a coupling between changes in math performance and refinement of fiber orientation, rather than between math changes and increases in fiber density. This pattern is striking when set against the developmental literature on NODDI, and our current findings when assessing baseline to endline changes in FA, ODI, and NDI across first grade. Across the first two decades of life, cross-sectional studies report that the steady, large-magnitude increases in FA observed during normative maturation are driven primarily by NDI, with a smaller and often inconsistent contribution from ODI (Genc et al., 2017; Mah et al., 2017). Our results indicate that the same biophysical drivers do not necessarily underlie short-timescale, learning-coupled microstructural change.

A “refinement, not proliferation” account fits the developmental context. By the time children enter first grade, the PVP tracts have already attained substantially adult-like microstructure (Vinci-Booher et al., 2022), implying that the foundational scaffold of neurites in these pathways is largely in place. What remains to develop, and what may be most responsive to ongoing experience, is the organizational refinement of those fibers, pruning of disorganized projections, tighter alignment of existing axons, and supporting glial reorganization (Genc et al., 2017; Zhang et al., 2012). Learning-related mΔODI may therefore index the structural signature of an architecture transitioning from “built” to “tuned” during the period when foundational math skills are consolidating for fluent first-grade math. Together, these findings point to a general principle: the slow accretion of new neurites and myelin that drives years-long development may not be how short-term, experience-dependent plasticity is implemented in early elementary-age white matter.

### Distributed, not window-specific, coupling between right-MdLF microstructure and math changes

We had hypothesized that one of three a priori windows — early (Sessions 1–3), ongoing (Sessions 3–10), or recent (Sessions 10–12) — would carry disproportionate weight in the right-MdLF–math coupling identified in Sections 4.3–4.4. No window did (pre-registered analysis with statistically necessary deviations). In both right-MDLF subtracts, no single-window model reached significance, and a comprehensive horse-race model including all three windows simultaneously also failed to identify any window as a significant predictor of total math gain. These findings indicate that the right-MDLF–math coupling is distributed: every month-to-month interval contributes incrementally, and no single month-slice of the academic year drives the relationship on its own.

This null result is informative for the methodological argument that has run through the present study. If learning-coupled white-matter change were concentrated in a discrete window of plasticity, then a sparsely sampled longitudinal design (e.g., a baseline + mid-year + endpoint scan) would be sufficient to capture the relevant signal, provided the timing of the scans was well chosen. Our results indicate the opposite: because the coupling is distributed across the year, sparse sampling would risk missing some contributing intervals while overweighting others, and the ΔFA–math relationship recovered in Section 4.3 is one that becomes statistically and theoretically tractable only when the full month-to-month time series is available. In this sense, the absence of a critical window is itself a positive empirical case for dense longitudinal neuroimaging in early elementary-age cohorts.

In sum, the first year of formal schooling refines the foundation for all subsequent academic learning, yet how the developing brain supports fundamental learning has been out of reach of conventional neuroimaging designs that sample too sparsely to resolve the dynamics of change. By following nine first-grade children with twelve monthly diffusion MRI scans and matched behavioral assessments, we show that brain and behavior are coupled on month-to-month timescales: behavioral trajectories were nonlinear, and concurrent fluctuations in the right middle longitudinal fasciculus tracked concurrent changes in math learning, driven by orientation refinement of existing fibers rather than neurite proliferation. The architecture of foundational learning is dynamic, distributed, and mechanistically distinct from the developmental trajectories we know — and seeing it requires watching it as it happens.

## Method

### Participants

Participants were recruited from the Nashville metropolitan area via social media, school-distributed flyers, and word-of-mouth. All participants were 6-year-old, right-handed, native English speakers who had completed at least one year of kindergarten and were entering first grade. Inclusion criteria required normal or corrected-to-normal vision and no history of neurological, developmental, or learning disorder diagnoses.

A total of 16 children were enrolled in July 2024. Six participants withdrew within the first six visits due to scheduling conflicts (N=4) or excessive head motion and lack of compliance (N=2). Of the remaining 10 participants who completed the full year of study, one was subsequently excluded from analysis due to abnormal brain structure. The study protocol was approved by the Vanderbilt University Institutional Review Board (IRB). Written informed consent and assent were obtained from parents or legal guardians and participants. All participants received monetary compensation for their time, and children were provided with a book and a toy of their choice at each visit.

### Study Design and Procedure

This study utilized a dense longitudinal neuroimaging design (Vinci-Booher et al., 2025). Participants completed 12 monthly visits over a 12-month period, beginning in the summer prior to first grade (e.g., July 2024) and concluding at the end of the academic year (e.g., June 2025). Each visit lasted approximately 2 hours, comprising a 60-minute MRI session followed immediately by a 60-minute behavioral assessment. During the first visit, participants completed a mock scanning session to acclimate to the MRI environment. During actual scans, foam padding was used to minimize head motion, and children watched a movie of their choice to promote stillness.

### MRI Data Acquisition

Imaging data were collected on a Philips 3T dStream™ Ingenia scanner equipped with a 32-channel head coil at the Vanderbilt University Institute for Imaging Science (VUIIS). High-resolution T1-weighted (1.0 mm isometric resolution; TR = ∼8.9 ms; TE = ∼4.6 ms; flip angle = 8°) and T2-weighted (1.0 mm isometric resolution; TR = ∼2500 ms; TE = ∼251.6 ms; flip angle = 90°) anatomical scans were acquired. Diffusion-weighted imaging (dMRI) was acquired using a multi-shell spin-echo echo-planar imaging (SE-EPI) sequence with a posterior-to-anterior phase-encoding direction. Scan parameters included: TR = 3200 ms, TE = 80 ms, flip angle = 90°, and a 2.5 mm isotropic voxel size with an in-plane acceleration factor of 2 and a multiband factor of 2. The sampling scheme consisted of 99 total volumes: 30 directions at b = 1000 s/mm^2^, 60 directions at b = 2000 s/mm^2^, and 9 interspersed non-diffusion-weighted (b ≈ 0 s/mm^2^) volumes. A separate reverse phase-encoded (anterior-to-posterior) scan of 6 non-diffusion-weighted (b ≈ 0 s/mm^2^) volumes was acquired to correct for susceptibility-induced distortion.

### Anatomical template construction

For each participant, a high-quality anatomical reference was constructed from the fMRIPrep longitudinal pipeline, which co-registered and averaged all usable T1-weighted scans across session into a single subject-level T1 template (Esteban et al., 2019; Reuter et al., 2012). A corresponding subject-level T2-weighted image was similarly derived, which was used for improved pial surface estimation. All T1 and T2 images were visually inspected using ITK-SNAP; only those of high quality were selected to create the subject templates. The median number of images used was 4 for T1 and 7 for T2. The T1 template was aligned to ACPC orientation using the HCP alignment tool (Glasser et al., 2013; Jenkinson et al., 2002), producing a 1 mm isotropic T1 reference volume (182 × 218 × 182 voxels) that served as the fixed coordinate frame for all subsequent processing.

### Diffusion preprocessing and cross-session alignment

Diffusion-weighted images for each session were preprocessed using a customized MRtrix3-based pipeline (Tournier et al., 2019). Preprocessing steps included MP-PCA denoising, Gibbs ringing removal, eddy current and motion correction via FSL eddy with slice-to-volume outlier replacement, EPI susceptibility distortion correction via topup using reverse phase-encoded b0 volumes, ANTs N4 bias field correction, and Rician noise estimation. Each session’s preprocessed DWI was then registered to the participant’s ACPC-aligned T1 template via boundary-based registration (BBR) computed with FSL, to ensure alignment across sessions.

### Generation of participant-specific tractograms

Anatomically constrained tractography (ACT) was performed per session in MRtrix3 with the iFOD2 probabilistic algorithm, generating 2,000,000 streamlines per session (step size 1 mm, maximum curvature 50°, length range 25-220mm) (Smith et al., 2012). Streamlines were seeded from the gray–white matter interface defined from a five-tissue-type anatomical segmentation of the participant’s T1 template. To support longitudinal analyses, a single participant-specific bundle template was then defined: White matter streamline classification was performed once per participant using a single reference session’s tractogram (session 1) in combination with a FreeSurfer segmentation (also computed once per participant from the subject-level T1 and T2 templates). Bundle labels were assigned via automated white matter segmentation and refined with iterative outlier removal (centroid SD = 4, length SD = 4, 5 iterations) (Bullock et al., 2022b; O’Donnell & Westin, 2007).

### Microstructural measurement at each session

Diffusion tensor fitting and multi-shell multi-tissue constrained spherical deconvolution (CSD) were computed per session to derive fiber orientation distribution and tensor-based microstructural maps (Basser et al., 1994; Tournier et al., 2007). For each session, microstructural metrics (FA, NDI, ODI) were sampled along these participant-specific bundle definitions following a tract profile approach (200 equidistant nodes per bundle; Yeatman, Dougherty, Myall, et al., 2012a).

### Behavioral assessments

Following each MRI scan, trained researchers administered math and reading assessments using FastBridge. Multiple forms of the assessment were used to allow for repeated monthly testing without practice effects. The math battery included Numeral Identification (naming numbers), Number Sequencing (answering sequence questions), Decomposing (part-part-whole problems), and Place Value (identifying numerals from base-10 blocks). The reading battery included Word Segmenting (segment words into individual sounds), Nonsense Words (reading pseudo-words), Sight Words (high-frequency words), and CBMreading (reading a passage aloud). Study sessions were video recorded and reviewed by trained research assistants for quality assurance. Math and reading were quantified as vertically-scaled composites derived from FastBridge earlyMath and earlyReading subtest raw scores. Each subtest was z-transformed against the G1_Fall mean and SD of our sample to place observations from all sessions on a common scale, then combined into a weighted composite using FastBridge’s published subtest weights. Composites were standardized to *M* = 50, *SD* = 15 using the G1 mean and SD of our sample. Critically, all standardization parameters were computed once and held fixed across all 12 sessions, so the value 50 represents the same anchored ability level at every time point (the mean G1 weighted-composite in our sample). A within-child increase therefore reflects performance growth on a common developmental scale rather than movement against a shifting age norm.

### Statistical analysis

All analyses were preregistered (https://osf.io/97ybe). Several adjustments to the preregistered plan were made based on diagnostic evaluation of the data; all deviations and post hoc analyses are documented below. All statistical analyses and visualizations were performed in Python 3.12 on macOS, using *NumPy* and *pandas* for data manipulation, *SciPy* for classical statistical tests and nonlinear curve fitting, *statsmodels* for regression modeling (including OLS with cluster-robust standard errors) and false discovery rate correction, and *Matplotlib* and *seaborn* for figure generation. Code for all statistical analyses will be made available on Github.

### Longitudinal trajectory modeling (Hypothesis 1)

To characterize developmental trajectories over the first-grade year, we fitted separate linear regression models for each participant to extract individual slopes (rates of change) for each outcome variable: composite math scores, composite reading scores, and mean FA for each of the 8 PVP tracts and the comparison tract. To test whether there was consistent development at the group level, we evaluated whether the mean slope across participants was significantly different from zero. The normality of the slope distributions was assessed using the Shapiro-Wilk test. For normally distributed slopes, a one-sample *t*-test was used; for non-normally distributed slopes, the non-parametric Wilcoxon signed-rank test was applied (pre-registered).

We further characterized the nature of behavioral change using two complementary approaches. First, we directly compared baseline (session 1) and endline (session 12) scores using paired-samples *t*-tests (or Wilcoxon signed-rank tests, as appropriate) to estimate the magnitude of overall changes from the start to the end of the school year. While the linear slope analysis tests whether improvement is consistent across sessions, the paired comparison offers a more intuitive and interpretable metric of total annual change (post hoc). Second, visual inspection of the raw trajectories suggested that linear models may not adequately capture the shape of learning across the year — reading trajectories appeared to rise rapidly before leveling off, while math trajectories showed an initial period of improvement followed by decline. Motivated by these observations, we fitted nonlinear growth models to the behavioral data: a sigmoid function for reading and a quadratic function for math. Together, these complementary analyses allow us to quantify both the amount and the shape of academic learning over the year (post hoc).

### Month-to-month brain–learning relations (Hypothesis 2)

To test our second hypothesis that microstructural properties would prospectively predict subsequent monthly changes in reading and math on a month-to-month basis, the preregistered analysis specified a dynamic panel linear mixed-effects model (LMM) of the form: Beh_Score(t) ∼ 1 + Beh_Score(t−1) + FA(t) + FA(t−1) + (1 | Subject). Two diagnostic issues necessitated modification of this plan. First, behavioral scores exhibited extremely high temporal autocorrelation (reading: Pearson’s *r* = 0.91, *p* < .0001; math: *r* = 0.80, *p* < .0001), indicating that a lagged dependent variable would absorb nearly all outcome variance and leave little to be explained by the white matter predictors. Second, LMMs with a random intercept for the subject showed persistent convergence failures, indicating that the model was overparameterized for the sample.

To address these issues, we adopted a complementary approach (pre-registered analysis with statistically necessary deviations) to the preregistered analysis. First, the dependent variable was redefined as a change score (Beh_Change = Beh_t − Beh_t−1), directly capturing month-to-month change in academic abilities. Second, we replaced the LMM with Ordinary Least Squares (OLS) regression with standard errors clustered by subject, which yields consistent fixed-effect estimates while correcting for within-subject dependence, and is well-established for small-sample longitudinal designs where LMMs may fail to converge due to insufficient data to identify random-effect variance components (Cameron & Miller, 2015; McNeish & Stapleton, 2016). We specified the primary Level-and-Change model, which regresses the session-to-session change in behavioral scores on both the baseline level of FA (FA_Lag1) and the concurrent change in FA (FA_Change). This formulation allows us to estimate the association between white matter change and behavioral change while controlling for baseline white matter status, providing a test of whether concurrent changes in FA are associated with concurrent learning changes, beyond what would be predicted by baseline FA alone. The model is specified for each tract as: Beh_Change

∼ FA_Lag1 + FA_Change + (clustered SE by Subject), where FA_Change = FA_t − FA_t−1 and FA_Lag1 = FA at time t−1. These models were fitted separately for each of the 8 tracts and for each academic domain (math and reading), yielding 16 tests per model specification (i.e., 8 PVP tracts x 2 academic domains). To control false discoveries while preserving sensitivity for our pre-registered hypotheses, tract-wise tests were assigned to three pre-defined hypothesis families — confirmatory (bilateral pArc for reading, bilateral TPC for math), exploratory (bilateral MDLFang and MDLFspl), post hoc (bilateral Arc, SLF1&2, SLF3, VOF, and the anterior frontal corpus callosum), and the Benjamini–Hochberg FDR procedure was applied separately within each family × domain × predictor combination, so that the larger exploratory and post hoc sets would not dilute power for the pre-registered confirmatory contrasts (FDR *q* < .05).

### Dissociation testing (Hypothesis 3)

For our Hypothesis 3, we predicted a functional double dissociation among the PVP sub-tracts: the pArc would be preferentially coupled to reading changes, while the TP-SPL would be preferentially coupled to math changes. The preregistered analysis proposed a hierarchical approach using linear mixed models (LMMs) with each participant’s individual learning slope as the dependent variable. This approach involved two steps: first, testing for a single dissociation within each tract via a two-way interaction between the brain metric (FA) and cognitive domain (math *vs.* reading); and second, testing for a double dissociation across tracts via a three-way FA × Tract × Domain interaction.

Given the convergence issues with LMMs described above, and our shift to a change-score framework, we adapted this analysis to an OLS regression with standard errors clustered by subject. Specifically, for tracts that showed a significant brain–behavior association in the primary month-to-month analysis, we tested for domain specificity by fitting the following model: Beh_Change ∼ FA_Change × Domain + (clustered SE by Subject), where FA_Change is the session-to-session change in fractional anisotropy and Domain indicates whether the outcome is change in math or reading scores. A significant FA_Change × Domain interaction would indicate that the tract’s coupling to learning differs between the two domains, providing evidence for a single dissociation. This approach tests the same conceptual question as the preregistered analysis — whether a given tract is selectively coupled to one cognitive domain — but in a framework consistent with our other change-score analyses.

### Secondary analysis 1: Microstructural specificity (NODDI)

As preregistered, this analysis was conducted only for tract–domain combinations that showed significant FA–learning relationships in the primary analyses to further interrogate microstructural changes underlying the FA change. For each such combination, we fitted identical OLS models with clustered standard errors, replacing FA with NDI and ODI.

### Secondary analysis 2: Critical predictive windows

To examine the critical predictive windows, there are necessary deviations from preregistration. The preregistered analysis modeled end-of-year scores as a function of FA at specific single time points (t1, t6, t9), controlling for initial scores. However, we revised this analysis to adopt a more dynamic framework consistent with the change-score approach used in the primary analyses. The outcome variable was redefined as total change over the year (Beh_Change_t1−t12), and FA change was partitioned into three non-overlapping temporal windows reflecting distinct phases of plasticity. Time points for windows (t1, t3, t10, t12) were selected based on three considerations. First, these sessions had complete data across all participants, maximizing statistical power. Second, the windows align with the academic calendar of the first year of formal schooling: the initial window (t1–t3; FA_t3 - FA_t1) captures the first few months of classroom exposure, a period often associated with rapid adaptation to novel learning demands; the ongoing window (t3–t10; FA_t10 - FA_t3) spans the main instructional period, during which the bulk of academic content is delivered; and the recent window (t10–t12; FA_t12- FA_t10) captures the end-of-year and early summer period, reflecting consolidation and retention following peak instruction. Third, this partition aligns with our observed behavioral trajectories: the quadratic fit of math scores peaked at approximately session 6.3 (**Figure 2**), which falls within the ongoing window, allowing us to test whether FA change during this period of peak behavioral growth is selectively predictive of overall behavioral change. Together, this framework enables us to test which temporal phase of white matter plasticity — initial acceleration, sustained development, or late consolidation — most strongly corresponds to year-long learning.

Specifically, for tracts showing significant FA–behavior relationships, we fitted four models:

Model A (Initial acceleration):

Beh_Change(t1–t12) ∼ 1 + FA_Change_Initial(t1–t3)

Model B (Ongoing development):

Beh_Change(t1–t12) ∼ 1 + FA_Change_Ongoing(t3–t10)

Model C (Recent consolidation):

Beh_Change(t1–t12) ∼ 1 + FA_Change_Recent(t10–t12)

Model D (Horse race):

Beh_Change(t1–t12) ∼ 1 + FA_Change_Initial + FA_Change_Ongoing + FA_Change_Recent

Models A–C tested each window in isolation, while Model D entered all three plasticity windows simultaneously to identify which, if any, uniquely predicted total learning changes. *P*-values were FDR-corrected across models.

